# Increased likelihood of heat-induced large wildfires in the Mediterranean Basin

**DOI:** 10.1101/2020.01.09.896878

**Authors:** J Ruffault, T Curt, V Moron, RM Trigo, F Mouillot, N Koutsias, F Pimont, NK Martin-StPaul, R Barbero, J-L Dupuy, A Russo, C Belhadj-Kheder

## Abstract

Wildfire activity is expected to increase across the Mediterranean Basin because of climate change. However, the effects of future climate changes on the combinations of atmospheric conditions that promote large wildfires remain largely unknown. Using a fire-weather based classification of wildfires, we show that future climate scenarios point to an increase in the frequency and severity of two heat-induced fire-weather types that have been responsible for a majority of record-breaking wildfire events. Heat-induced fire-weather types are characterized by compound dry warm conditions and occur in the summer during heatwaves, either under moderate (*sudden heatwave* type) or intense (*hot drought* type) drought. Heat-induced fire weather is projected to increase in frequency by 14% by the end of the century (2071-2100) under the RCP4.5 scenario, and by 30% under the RCP8.5. These findings suggest that the frequency and extent of large wildfires will increase throughout the Mediterranean Basin, with far-reaching impacts.

---

Climate is the major driver of wildfires at the regional scale and an increase in climate-induced wildfire activity has been documented in a number of ecosystems over the past decades^1,2^. In contrast, fire activity has been declining in most Euro-Mediterranean countries owing to management and suppression measures undertaken since the 1980s^3^. Nevertheless, recent extreme wildfire events, including those that occurred in 2016 in France^4^, 2017 in Spain and Portugal^5^ and 2018 in Greece^6^ have highlighted the limits of fire suppression capabilities under exceptional weather conditions. While previous showed that wildfire activity is expected to increase across the Mediterranean Basin because of climate change^7,8^, how the combinations of atmospheric conditions that promote large wildfires will be affected remain largely unknown and unquantified.

In temperate mid-latitude ecosystems, a wide range of meteorological fields (precipitation, temperature, relative humidity and wind speed) influence the spread of wildfires on multiple timescales. Moisture deficits over days to months increase fuel aridity^1,9^, which interact with meteorological conditions during the wildfire to govern its behaviour.^10,11^. Most of the largest wildfires occur when certain extreme weather events overlap^12^. For instance, the combination of extreme drought with extreme wind or heatwaves have both been identified as crucial factors in wood-fueled crown wildfires in Mediterranean forests and shrublands^13,14^‥ Evaluating how and to what extent the expected warming and drying of the climate in the Mediterranean Basin will affect the risk of large wildfires is crucial for mitigation and adaptation planning^15^

The holistic approach adopted in this study was to investigate fire-weather relationships at the continental level by defining five fire-weather types (FWTs) based on different combinations of climate variables that are critical for wildfire spread. We have analyzed >17,000 records of wildfires (> 30 ha) in four countries (France, Greece, Portugal and Tunisia) covering most of the diverse biogeographic and climatic conditions found in the Mediterranean Basin (Supplementary Figs. 1–3).

The set of weather conditions under which wildfires occur (the “wildfire niche”) was described by five variables that describe different levels of fuel aridity and the synchronous, short-term conditions that control the spread of wildfires in Mediterranean ecosystems (see *Methods*). The variables are the mean temperature, wind speed, relative humidity, and two measures of fuel aridity, the duff moisture code (DMC) and the drought code (DC)^16^, which account for the level of drought in the preceding weeks and months, respectively. Each wildfire in the database was related to the daily values of the climate variables observed at the closest gridcell. To facilitate interpretation, the observations were expressed in terms of two orthogonal linear combinations of the climate variables by principal component analysis (PCA) (Fig. 1).

**Fig. 1.**
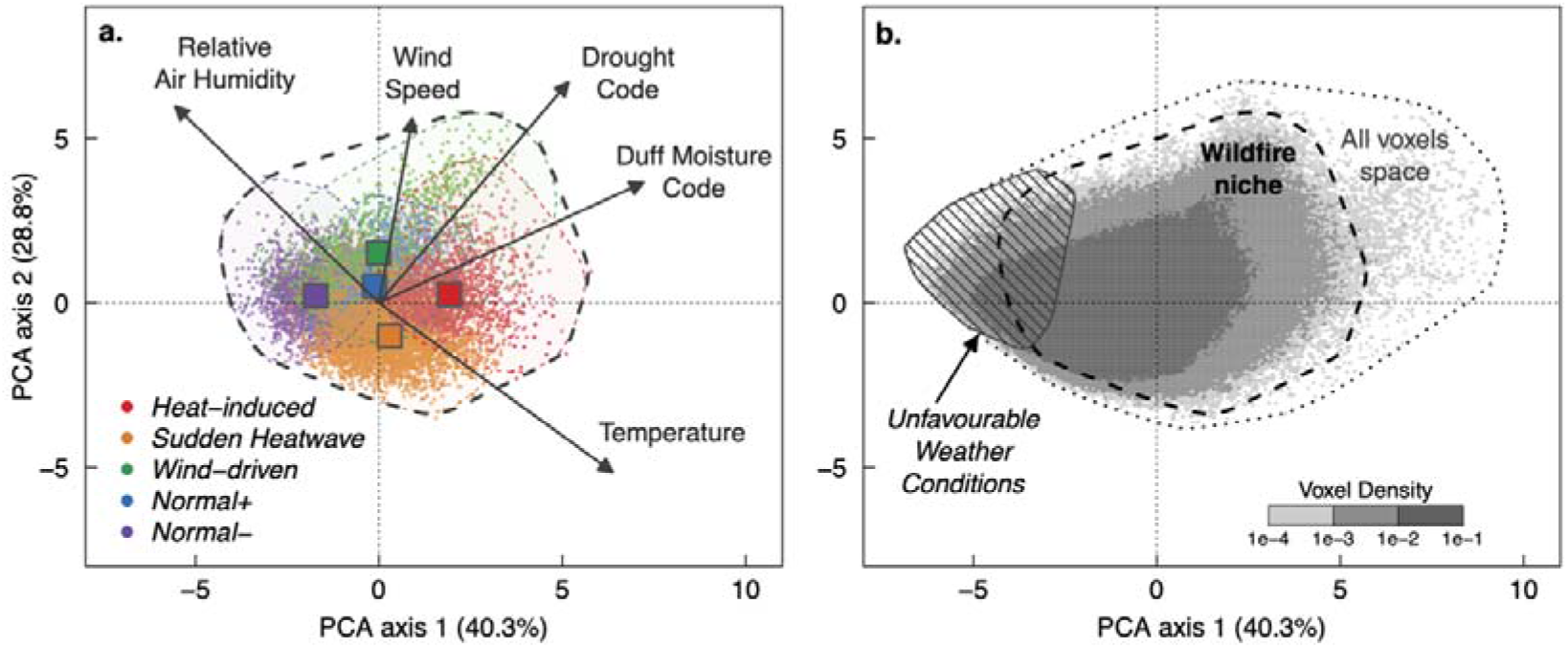
**Fire-weather types (FWTs) in the wildfire niche** shown in terms of the first two orthogonal combinations of wildfire predictors determined by principal component analysis (PCA) of five daily climate variables driving wildfires. Wildfire records (> 30 *ha*) were extracted for the period 1985– 2015 from four countries covering most of the biogeographical conditions found in the Mediterranean Basin (France, Greece, Portugal and Tunisia). **a.** Local daily values of the climate variables associated with each wildfire (colored dots) and centroids (colored squares) for each FWT. **b**. The wildfire niche is the atmospheric space that contains all the wildfires (> 30 ha) in the dataset, the *All Voxels* space covers all days and grid cells (including non-fire voxels) in the summer fire season. The *Unfavourable* weather conditions correspond to voxels in which wildfires are unlikely occur because the weather conditions are moister and cooler than those observed in the wildfire niche.

The wildfire niche in the Mediterranean Basin encompasses a broad range of conditions (Fig. 1a). This is consistent with the substantial disparities in fire-climate relationships that have been reported across the Mediterranean Basin^11,14^. The wildfires were classified into five fire-weather types (FWTs) by cluster analysis of the climate variables^14^ (Fig. 1a, details in *Methods*). The probability of a wildfire spreading over larger areas under each FWT was quantified by the large fire risk ratio (LFRR), defined as the probability of a wildfire spreading to a certain size under a given FWT divided by the corresponding probability under the other FWTs (see *Methods*). The FWTs fall into three broad categories according to their main characteristics.

The baseline category consists of two FWTs, *Normal*− and *Normal*+, characterized by little or no change in temperature, humidity or wind speed on the day of the wildfire compared with the preceding or following days (Fig. 2a). The main difference between *Normal*− and *Normal*+ is that the former is significantly moister than the other FWTs, including *Normal*+ (Fig. 2a), but remains in the average of observed summer conditions, except in Tunisia (Supplementary Figs. 5 and 6). Accordingly, *Normal*− wildfires are more frequent in the northern part of the Mediterranean Basin (Fig. 2c) and tend to occur sooner during the fire season than *Normal*+ wildfires do (Supplementary Fig. S4). The LFRRs are significantly lower for baseline FWTs than for the other FWTs (Fig. 2b and Supplementary Table S2). In particular, *Normal*− is associated to the lowest risk of large wildfires, accounting for 19 % of fires but just 9 % of the total burned area (Fig. 2b and Supplementary Table S2).

**Fig. 2.**
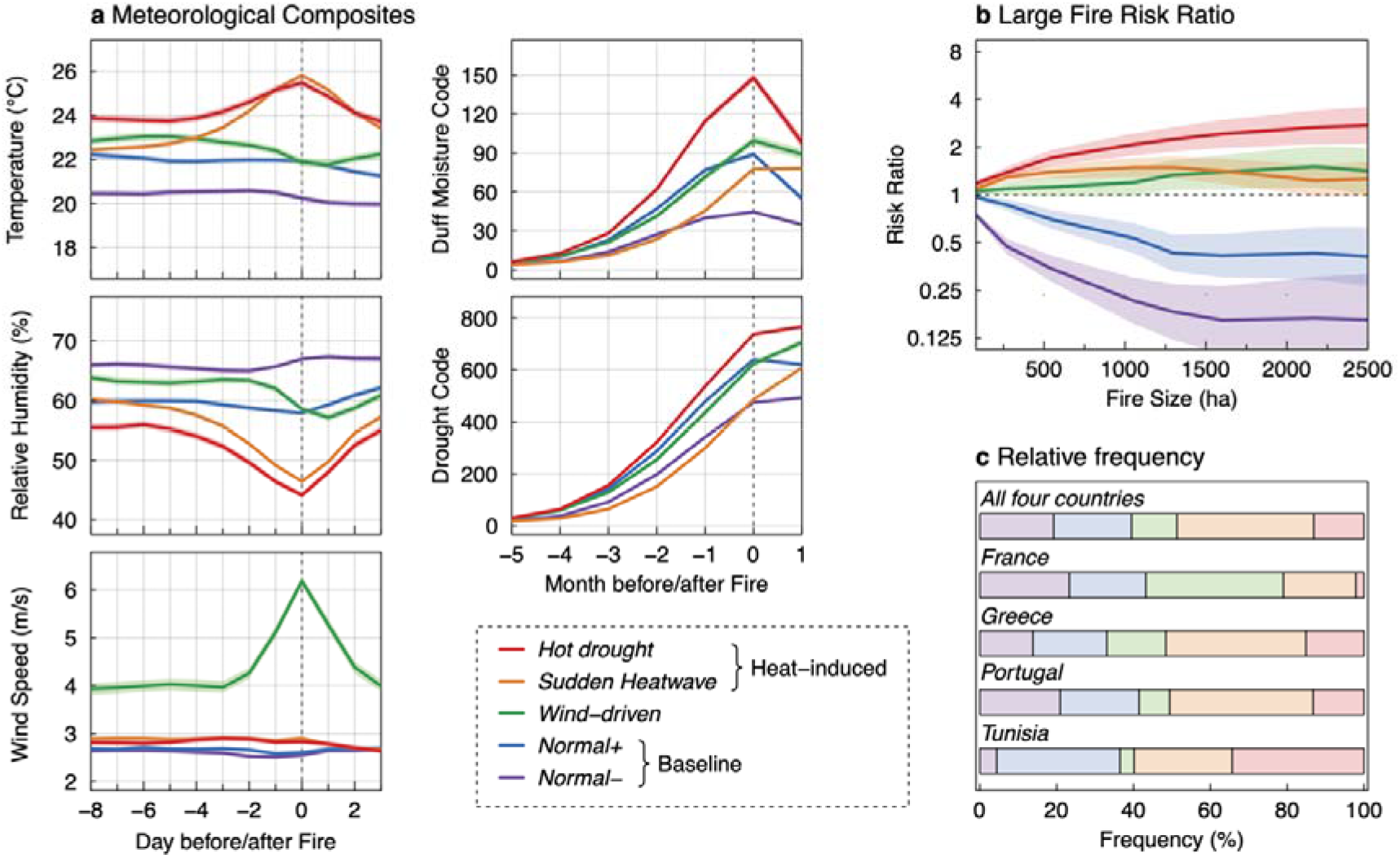
Fire weather type (FWT) characteristics. **a.** Lead lag climate variable composites on daily (for temperature, mean relative humidity and mean wind speed) and monthly (for drought code and duff moisture code) timescales. For each FWT, the mean values (lines) and the 95% confidence interval around the mean (coloured areas) are reported. The dashed grey vertical lines indicate the day (month) of a fire. **b.** The large fire risk ratio (LFFR) for each FWT: mean values (lines) and 95% confidence intervals (coloured areas). The LFFR is the ratio of the probability of a wildfire reaching a certain size under a given FWT divided by the corresponding probability under the other FWTs. **c.** Relative frequency of the FWTs in the four Mediterranean countries investigated here.

The two heat-induced types of fire weather, *Sudden Heatwave* and *Hot Drought*, are both characterized by drought conditions (higher DC and DMC than the averaged summer conditions) with higher temperature and lower humidity on the day of the wildfire compared with the preceding and following days (Fig. 2a, and Supplementary Figs. 5 and 6). The temperature and relative humidity on wildfire days under these FWTs are similar, but *Hot Drought* conditions are the driest of all, with higher levels of DC and DMC than all other FWTs (Fig. 2a). *Hot Drought* wildfires are thus more frequent in the southern part of the Mediterranean Basin (Fig. 2c) and tend to occur later in the fire season than *Sudden Heatwave* wildfires do (Supplementary Fig. 4). While both heat-induced FWTs are associated with higher LFRRs than the background FWTs are, the risk of very large wildfires (> 1,500 ha) is particularly high under *Hot Drought* conditions.

The final FWT identified in this analysis accounts for wildfires induced by strong winds on the day of the wildfire and relatively dry conditions (Fig. 2a). *Wind-driven* wildfires occur mainly in coastal areas, notably Southern France and Greece (Fig. 2c), where wildfires are often driven by continental northerly synoptic winds respectively known as the Mistral^10^ and the Etesian winds^17^. *Wind-Driven* conditions are associated with higher LFRRs than are the background FWTs.

Our fire-weather based classification highlights the fact that large wildfires occur mostly when short-term meteorological extremes combine with long-term summer drought. Strong surface winds coupled with low relative humidity and drought conditions are known to favor large wildfires worldwide^10,13^. However, the most extreme wildfires in the Mediterranean Basin occur under heat-induced *Hot Drought* conditions, when drought and temperatures are the highest. This conclusion matches previous observations on the prevalence of wildfires and extreme behaviours and intensities ^5,18^ on particularly hot days or during heatwaves in the Mediterranean Basin^5,14,19^. The mechanisms underlying these wildfires remain unclear but the fast desiccation of live fuel due to warming◻induced increases in the vapor pressure deficit probably contributes. Besides, it must be noted that the levels of DC associated with the FWTs (Supplementary Figs. S5 and S6) confirm indirectly that seasonal fire activity depends mainly on fuel aridity in the moister northern part of the Mediterranean Basin ^20–22^, and on meteorological conditions associated with influxes of dry, hot air in the drier southern part.

Fire-weather types offer new insights into the climate-induced changes in wildfire potential, because they disentangle the underlying multi-scale combinations of atmospheric conditions that are conducive to wildfires. The fire-weather classification was therefore applied to summer conditions (June to September) throughout the Mediterranean Basin (independently of the occurrence of wildfires) using simulations from eight climate models produced as part of the EURO-CORDEX initiative^23^ for RCP4.5 and RCP8.5 emission scenarios (Supplementary Table S1). To extend the fire-weather classification to summer weather conditions during which wildfire are unlikely to occur, *i.e*. when fuel moisture prevents significant wildfire spread^9^, the additional type *Unfavourable* was added (left side of the principal component subspace, Fig. 1b).

This analysis reveals large and robust shifts in the frequency of the FWTs towards those associated with heat-induced wildfires by the end of the century (Fig. 3, right columns). The results are described here for the 2071-2100 period under the two concentration pathways because the mid-century (2031-2060) FWT frequencies under the two scenarios and both similar to the end-century frequencies under the RCP4.5 scenario (Fig. 3 and Supplementary Fig. S7). The relative frequency of heat-induced FWTs will increase on average by 14 % by the end of 21^st^ century under the RCP4.5 scenario and by 30 % under RCP8.5, with good agreement between the trends projected with the eight climate models (Supplementary Fig. S8). The projected changes in the frequencies of the FWTs differ markedly either side of 40^th^ parallel (Fig. 3). On the northern side, *Unfavourable* and *Normal*− conditions will become less common and *Normal*+, *Sudden Heatwave* and, to a lesser extent, *Hot-Drought* FWTs will increase in frequency. In the drier southern part (Supplementary Fig. S3), our results show that *Hot-Drought* conditions will increase in frequency at the expense of *Normal*− and *Normal*+ FWTs. Our study also demonstrates an increase in the intensity of *Hot-drought* and *Sudden Heatwave* (Supplementary Fig. S9) throughout the Mediterranean Basin (Supplementary Fig. S10). Uncertainty remains, however, as to the extrapolation of FWTs outside the current wildfire niche (Right side of the principal component subspace, Fig. 1b). In the present study, these voxels were related to the closest FWT centroid, while being aware that this hypothesis (conservative) might hide other fire-weather conditions that are not captured by the FWTs.

**Fig. 3.**
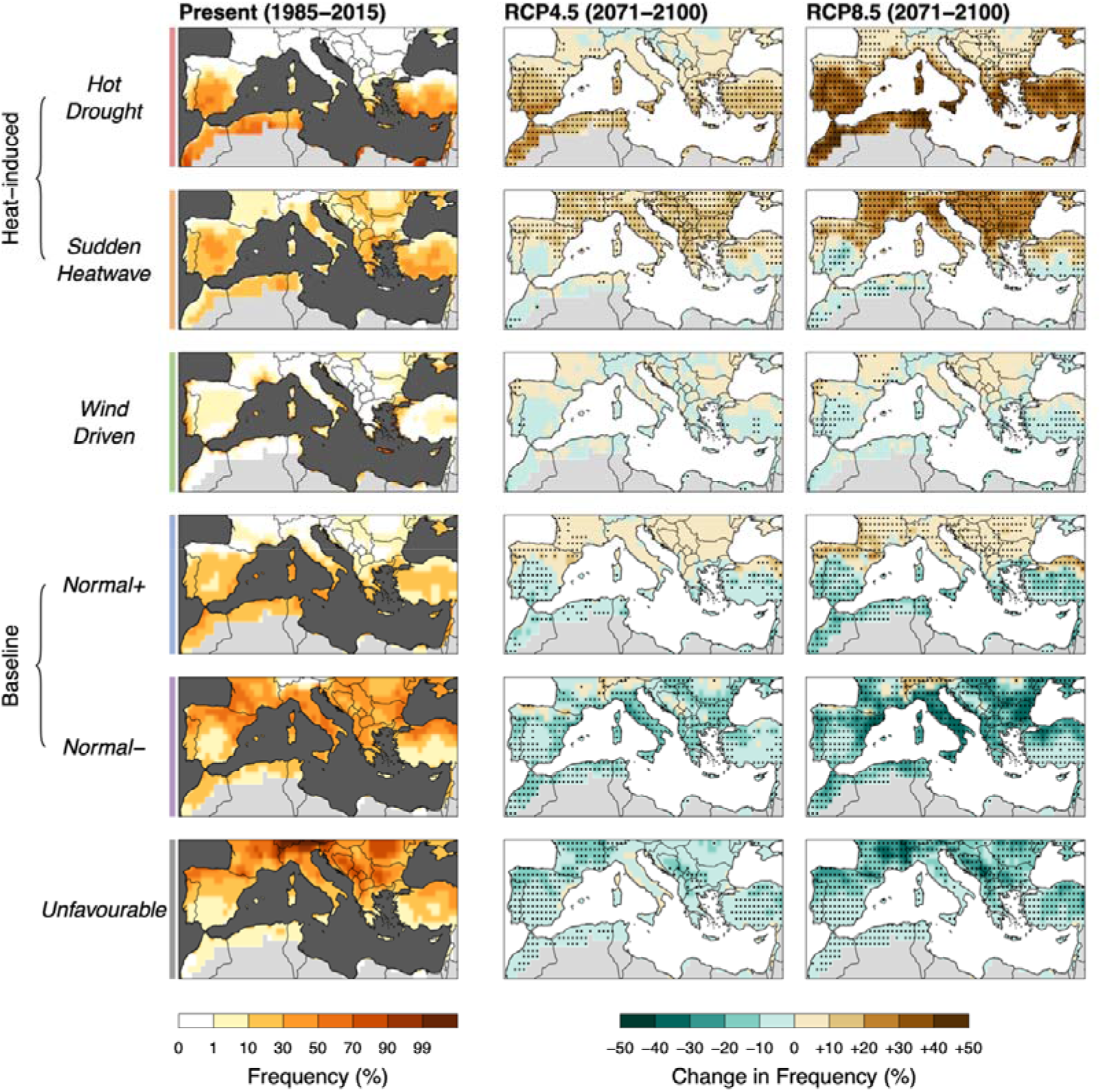
Current and multi-model median changes in summer frequency of fire weather types (FWTs) in the Mediterranean basin. The changes in frequency at the end of the century (2071– 2100) are shown relative to the present period (1985-2015) for two emission scenarios (RCP4.5 and RCP8.5). The dots indicate grid cells where the change in FWT frequency was statistically significant (*p* < 0.05) in a majority of the models.

Assuming that the ratio between wildfire frequency and FWT frequency would remain the same (stationarity of the fire-FWT relationship), changes in FWT frequency are expected to increase the number of wildfires (>30 *ha*) by 91% in France, 29 % in Greece, 21% in Portugal and 30 % in Tunisia by the end of the century under the RCP8.5 scenario (Supplementary Figs. S11 and S12). However, fire-weather relationships are influenced by non-climatic factors in many ways^14,24^, and thus so are wildfire-FWT relationships. This is particularly important in the Mediterranean Basin, because vegetation cover, human settlement patterns and fire suppression measures vary widely and are liable in the coming decades^25^. Improvements in suppression capacities and fuel management may therefore moderate the conclusions drawn her on the increased likelihood of wildfires^26^. Likewise, in the most arid regions of southern Europe, the fuel load and fuel continuity may be insufficient to sustain more frequent wildfires, and there may instead be a shift in fire–climate–vegetation relationships^24^, which are not captured by this FWT approach. In contrast, some factors may amplify the FWT–fire relationship, such as increased fire ignition at wildland-urban interfaces^27^.

Compound dry-warm periods are projected to become more frequent and persistent in a warmer climate^28^ and future scenarios of fire danger consistently point to an increase in the frequency and intensity of heat-induced FWTs across the Mediterranean. This will very likely increase the frequency and extent of wildfires, provided fuel remains abundant. The ecological and socio-economic implications of these changes could be profound, especially in view of the other emerging and interconnected risks that are projected to affect the Mediterranean Basin in the coming decades^15^.To mitigate the risks associated with heat-induced wildfires, dedicated efforts should be made to address current knowledge gaps on fuel desiccation during hot droughts^29^ and the effects of fuel moisture content on wildfire behavior^30^.

## Methods

We extracted the location, date, and size of wildfires that occurred in Southern France, Greece, Portugal, and Tunisia (Supplementary Fig. 1) between 1985 and 2015 from national fire databases. To limit uncertainties related to the detection rate of the smallest wildfires and increase the fire-weather signal, wildfires smaller than 30-ha size were excluded. Only summer wildfires (from June to September) were analyzed because 84 % of all wildfires occurred during these months and accounted for more than 82 % of annual burnt area (including of all wildfire sizes, Supplementary Fig. S2). The four countries were selected for two main reasons: (i) the availability for each one of a validated database with daily information on the characteristics, including the location and size, of all fire events; and (ii) the fact that together, they cover most of the biogeographic and socio-economic conditions found in the Mediterranean Basin^25^ (Supplementary Fig. S2). For Southern France, fire statistics were extracted from the “Prométhée” database, managed by French forest services, and examined extensively in previous studies^31,32^. For Greece, fire data from before 1998 were obtained from the Greek forest service and from 2000 onwards, the data were obtained from the Greek fire brigade^33^, such that data for the years 1998 and 1999 are missing. While this combination of sources may have led to some minor inconstancies, the potential consequences on the results were assumed to be negligible, as year of occurrence was not a factor in the analysis. Fire statistics for Portugal were extracted from the Portuguese rural fire database, described in detail in ref.^34^. For Tunisia, fire statistics were obtained from the Tunisian fire database^35^ as compiled from the records of the Tunisian Directorate-General for Forests and curated from various remote sensing sources. Wildfires from 2011 to 2015 were ignored because of the disruption in fire activity resulting from the Arab spring in December 2010^35^. Fire statistics for the four studied databases are shown in Supplementary Figure 2.

The meteorological data used to determine wildfire variables in the studied period and as a bias correction reference for the climate simulations were taken from the ECMWF ERA-Interim reanalysis^36^ of data from 1979 to 2015 across the Mediterranean Basin (see studied region in Supplementary Fig. 1). We ignored gridcells that contained more than 90% of non-burnable area, as determined from the ESA CCI Land Cover maps for the year 2015 (ESA 2019) at a spatial resolution of 300 m and aggregated at the ERA-INTERIM reference grid cell level. Daily variables were obtained from a 0.75° resolution geographic grid. We used the daily 2-m air temperature, dew point temperature, total surface precipitation, surface pressure and 10-m wind speed. Relative humidity was calculated using the mean daily dew point temperature, surface pressure and the mean daily 2-m air temperature.

The DMC and DC subcomponents of the Fire Weather Index^16^ were respectively used as generic indicators of medium and long-term drought. The DMC and DC are calculated in theory from daily 24□h accumulated precipitation data, and the temperature, humidity, and wind speed at 12:00 local time. However, since obtaining modeled data at 12:00 local time was not possible, we used the daily mean temperature, mean humidity and accumulated precipitation. This may have biased the DC and DMC estimates, but time series calculated with the two variables are highly correlated ^37^. Furthermore, as these fire variables were mostly used for classification purposes, we do not expect this approximation to have a significant impact on the results of the study. Nonetheless, the DC and DMC values calculated here may differ from those computed with the usual 12:00 data.

Projections of fire variables for the current climate and future projections were obtained from two regional climate simulation programs involved in the fifth phase of the Coupled Model Intercomparison Project (CMIP5) and produced as part of the EURO-CORDEX initiative^23,38^. Both include the same eight General Circulation model(GCM)-Regional circulation model (RCM) pairs (involving three RCMs and five GCMs; Supplementary Table 1), but one follows the moderate RCP4.5 scenario while the other follows RCP8.5, the highest concentration pathway^39^. RCMs were selected based on the availability of daily values for mean temperature, relative humidity, wind speed and precipitation. For each simulation, data were extracted at a spatial resolution of 0.44° in latitude and longitude for the historical (1970–2005) and future (2006–2099) periods in the studied regions (shown in Supplementary Fig. 1). Spatial and seasonal biases in projected climate variables can be problematic for the calculation of fire danger indices ^40^ so the data were corrected using univariate and multivariate bias correction based on daily summaries from ECMWF ERA-Interim data for the overlapping period between ECMWF ERA-Interim and climate simulation(1980-2005). The Euro-CORDEX data were interpolated to the regular 0.75° resolution grid of the ERA-INTERIM database using nearest neighbors prior to any other transformation. Statistical corrections were applied on a monthly basis to account for seasonal variations in distributional differences. We using the quantile delta mapping method^41^ for univariate bias correction and the MBCn algorithm^40^ for the multivariate correction. As both methods yielded similar results (Supplementary Fig. S13), only the results for the univariate biais correction method were used.

We identified FWTs objectively by dynamic k-means clustering based on the values of the fire predictor variable associated with each wildfire record. This method partitions *m* multivariate observations into *k* clusters in which each observation is allocated to the cluster with the nearest mean. In practice, the numerical algorithm minimizes the sum over all clusters of the within-cluster sum of observation-to-centroid squared Euclidean distances. As with any other dynamic clustering method, the value of *k* has to be chosen beforehand. To optimize our understanding of fire-weather relationships at the continent level, *k* was chosen as a trade-off between three competing criteria namely, (i) that *k* should be large enough to discriminate between the maximum possible combinations of wildfire predictors, (ii) that *k* should be small enough for the results to be generalizable and to facilitate interpretation, and (iii) that the final set of clusters should provide a sound physical interpretation of fire-weather relationships. We therefore set the clustering algorithm to identify *k*=5 FWTs. The stability of the clusters was verified by rerunning the algorithm with different initial random seeds. Furthermore, since the wildfires were not evenly distributed between the studied countries (Supplementary Fig. S2 and Supplementary Table 2), we also tested the stability of the clusters against variations in the proportion of wildfires from each country. The same k-means analysis was performed on 1,000 bootstrap-resampled wildfire datasets, in which the number of wildfires from each country was proportional to the corresponding number of wildfire gridcells. The FWTs in the original and resampled datasets had similar characteristics (Supplementary Fig. 14), confirming the robustness of this approach.

The climate variables were compressed into orthogonal linear combinations using PCA. Only the first two principal components were retained, as they adequately described most (69%) of the variability of the data. All gridcell*day (voxel) combinations (i.e. not just fire*day combinations) were then plotted in PCA space for the four studied countries. Voxels located outside the wildfire niche (line around the fire voxels) in the top-right quadrant of principal component subspace (Fig. 1b) were considered extreme representatives of the closest-centroid FWT. In contrast, voxels located outside the wildfire niche in the bottom-left quadrant of the principal component subspace (Fig. 1b) were considered representative of *Unfavourable* weather conditions. This additional FWT consists of all the voxels from the *Normal*− FWT whose Euclidian distance to the centroid was greater than the 95 % confidence interval. For each GCM-RCM (Table S1), the change in FWT frequencies between the current period (1985–2015) and two future periods (2031–2060 and 2071–2100) were calculated under the RCP4.5 et RCP8.5 concentration pathways.

The FWTs were characterized using a lead-lag analysis of composite climate variables. As wildfires in the Mediterranean Basin are short, we focused mainly on pre-ignition conditions. Lead-lag composites were examined over two timescales to capture the seasonal and synoptic variability associated with fire occurrence. We used an 11-day window (from 8 days before to 2 days after the start of a fire) for the meteorological variables (mean temperature, relative humidity and wind speed) and a 7-month window (from 5 months before to 1 month after the start of a fire; usual calendar months) for the fuel aridity variables. To assess the differences between the FWTs and normal summer conditions, standardized anomalies were compiled for each country: a reference climate was estimated in each country and the standardized anomalies were calculated in comparison. The reference climates were determined by randomly selecting 10,000 non-fire gridcell*day voxels with the same location in space and time (in the annual cycle) as the fires. This allowed a fair comparison between fire and non-fire atmospheric conditions with the same spatial and temporal frames.

To assess whether wildfires spread preferentially under particular FWTs, we calculated the LFRR for each FWT. The LFRR of a FWT and a fire size S is the ratio of the probability of a fire reaching size S in FWT T divided by the probability of a fire reaching the same size in the other FWTs. According to this definition, a LFRR higher (alternatively lower) than 1 indicates that a fire of size S is more (alternatively less) likely in FWT T than in the others. Fire size risk ratios were calculated for eight fire sizes (30, 80, 271, 550, 1055,1295, 1600 and 2173 ha), corresponding respectively to the 1^st^, 50^th^, 80^th^, 90^th^, 95^th^, 96^th^, 97^th^ and 98^th^ percentiles of the data (Supplementary Table S2).

## Acknowledgments

This work is a contribution from the Labex OT-Med (ANR-11-LABEX-0061) funded by the “Investissements d’Avenir”, a French National Research Agency (ANR) program, through the A*Midex project (ANR-11-IDEX-0001-02). The fire data for Tunisia were provided by Direction Générale des Forêts (Tunis) and fully checked and corrected within the FUME FP7 EU project Contract Grant No. 243888. N. Koutsias received funding from the European Union’s Horizon 2020 Research and Innovation Program under the Marie Skłodowska-Curie Grant Agreement No. 705067.

## Author Contributions

J.R., T.C., V.M. and R.T. conceived the study. J.R. performed the analyses with the help of T.C, V.M, R.T and F.P. N.K, F.M., A.R and C.B.K. provided datasets. JR wrote the manuscript and T.C., V.M., R.T., F.M, N.K, N.K.M., F.P, R.B. and J.L.D. provided comments and feedback.

## Competing Interests

The authors declare no competing interests.

## Notes

#### Summary of Updates

Corrected affiliations

