## supplementary material for "Increased likelihood of heat-induced large wildfires in the Mediterranean Basin"

*[1]Institut Méditerranéen de Biodiversité et d'Ecologie marine et continentale (IMBE) Aix Marseille Université, CNRS, IRD, Avignon Université, France*
*[2] Current address : INRAE, Ecologie des Forêts Méditerranéennes (UR 629), Avignon, France*
*[3] INRAE, Mediterranean Ecosystems and Risks, Aix-en-Provence, France*
*[4] Aix Marseille University, CNRS, IRD, INRAE, Coll. de France, CEREGE, Aix-en-Provence, France*
*[5] Instituto Dom Luiz (IDL), Faculdade de Ciências, Universidade de Lisboa, Campo Grande, 1749–016, Lisboa, Portugal*
*[6] CEFE, Univ. Montpellier, CNRS, EPHE, IRD, univ. Paul Valery Montpellier 3, 1919 route de Mende, 34293 Montpellier Cedex 5, France.*
*[7] Department of Environmental Engineering, University of Patras, G. Seferi 2, GR-30100, Agrinio, Greece*
*[8] INRAE, Ecologie des Forêts Méditerranéennes (UR 629), Avignon, France*
*[9] Geomatic and Geosystems, University of Mannouba-Tunis, Tunis, Tunisia*

| ID | Forcing GCM | Run | RCM | Institution |
|---|---|---|---|---|
| 1 | CNRM-CM5 | r1i1p1 | RCA 4 | Swedish Meteorological and Hydrological Institute (SMHI) |
| 2 | ICHEC-EC-EARTH | r12i1p1 | RCA 4 | Swedish Meteorological and Hydrological Institute (SMHI) |
| 3 | MPI-ESM-LR | r1i1p1 | RCA 4 | Swedish Meteorological and Hydrological Institute (SMHI) |
| 4 | IPSL-CM5A-MR | r1i1p1 | RCA 4 | Swedish Meteorological and Hydrological Institute (SMHI) |
| 5 | MHOC-HadGEM2-ES-01 | r1i1p1 | RCA 4 | Swedish Meteorological and Hydrological Institute (SMHI) |
| 6 | ICHEC-EC-EARTH | r1i1p1 | RACMO 2.2 | Royal Netherlands Meteorological Institute (KNMI) |
| 7 | MHOC-HadGEM2-ES-01 | r1i1p1 | RACMO 2.2 | Royal Netherlands Meteorological Institute (KNMI) |
| 8 | ICHEC-EC-EARTH | r3i1p1 | HIRHAM 5 | Danish Meteorological Institute (DMI) |

**Table S1** EURO-CORDEX experiments used in this study. Each experiment includes one historical and two scenario (RCP4.5 and RCP8.5) runs; spanning the periods 1970-2005 and 2006-2099 respectively.

| | Fire weather Type | Final fire size (ha) | | | | | | | |
|---|---|---|---|---|---|---|---|---|---|
| | | 30 | 80 | 271 | 550 | 1055 | 1295 | 1600 | 2173 |
| All four countries | *Normal-* | 2571 | 1076 | 291 | 108 | 35 | 24 | 16 | 11 |
| | *Normal+* | 2765 | 1446 | 531 | 222 | 88 | 58 | 42 | 29 |
| | *Wind-driven* | 1624 | 911 | 380 | 194 | 103 | 90 | 71 | 50 |
| | *Sudden Heatwave* | 4909 | 2787 | 1245 | 649 | 337 | 269 | 196 | 121 |
| | *Hot drought* | 1815 | 1122 | 518 | 308 | 179 | 151 | 120 | 86 |
| France | *Normal-* | 225 | 90 | 34 | 16 | 10 | 8 | 8 | 6 |
| | *Normal+* | 206 | 128 | 54 | 24 | 13 | 13 | 12 | 10 |
| | *Wind-driven* | 378 | 236 | 111 | 64 | 37 | 31 | 24 | 16 |
| | *Sudden Heatwave* | 192 | 100 | 55 | 27 | 18 | 13 | 8 | 3 |
| | *Hot drought* | 24 | 12 | 4 | 4 | 3 | 2 | 2 | 2 |
| Greece | *Normal-* | 435 | 145 | 30 | 5 | 2 | 2 | 0 | - |
| | *Normal+* | 617 | 290 | 101 | 30 | 10 | 9 | 8 | 6 |
| | *Wind-driven* | 505 | 289 | 131 | 60 | 37 | 32 | 26 | 20 |
| | *Sudden Heatwave* | 1182 | 655 | 282 | 142 | 76 | 64 | 48 | 25 |
| | *Hot drought* | 502 | 295 | 116 | 71 | 36 | 31 | 25 | 19 |
| Portugal | *Normal-* | 1907 | 840 | 227 | 87 | 23 | 14 | 8 | 5 |
| | *Normal+* | 1903 | 1006 | 372 | 165 | 63 | 34 | 21 | 13 |
| | *Wind-driven* | 737 | 384 | 137 | 69 | 28 | 26 | 19 | 14 |
| | *Sudden Heatwave* | 3504 | 2011 | 901 | 475 | 242 | 191 | 139 | 92 |
| | *Hot drought* | 1243 | 783 | 389 | 228 | 138 | 116 | 91 | 63 |
| Tunisia | *Normal-* | 4 | 1 | 0 | 0 | 0 | 0 | 0 | 0 |
| | *Normal+* | 39 | 22 | 4 | 3 | 2 | 2 | 1 | 0 |
| | *Wind-driven* | 4 | 2 | 1 | 1 | 1 | 1 | 1 | 0 |
| | *Sudden Heatwave* | 31 | 21 | 7 | 5 | 1 | 1 | 1 | 1 |
| | *Hot drought* | 46 | 32 | 9 | 5 | 2 | 2 | 2 | 2 |

**Table S2** Wildfire size distribution according to each fire weather type (FWT) for the entire wildfire dataset and for each of the country examined.

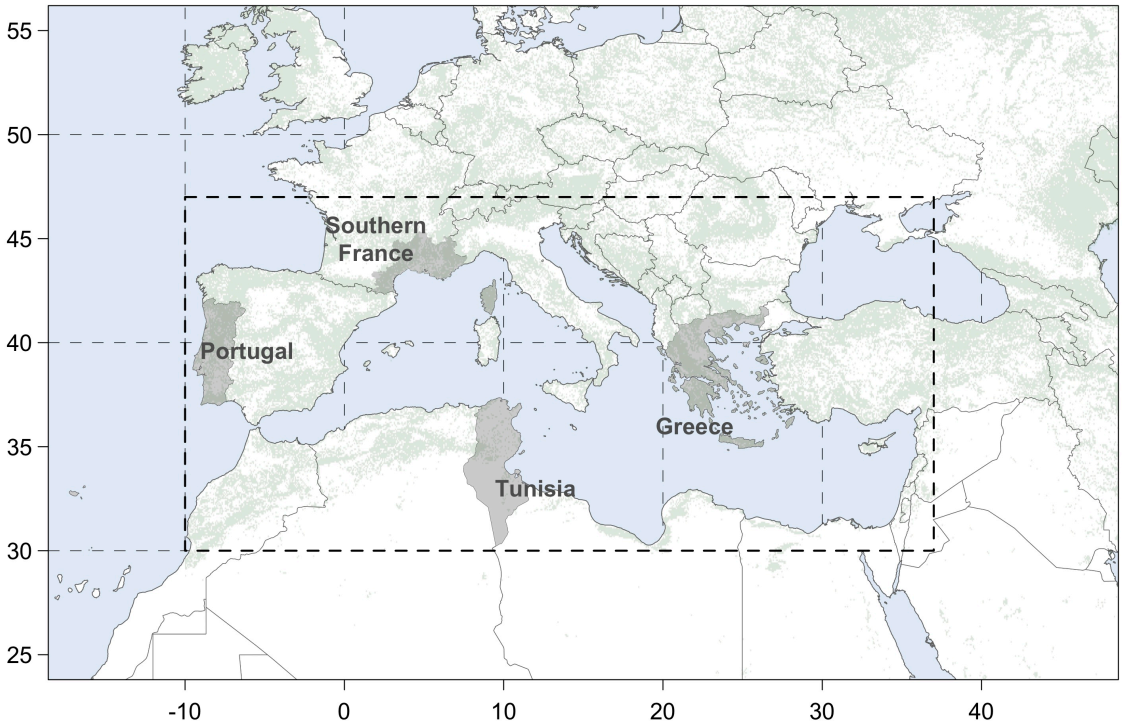


**Fig. S1** Location of the four countries for which wildfire records were examined (Southern France, Greece, Portugal, and Tunisia) within the Mediterranean geographic domain considered in this study (dashed rectangle).

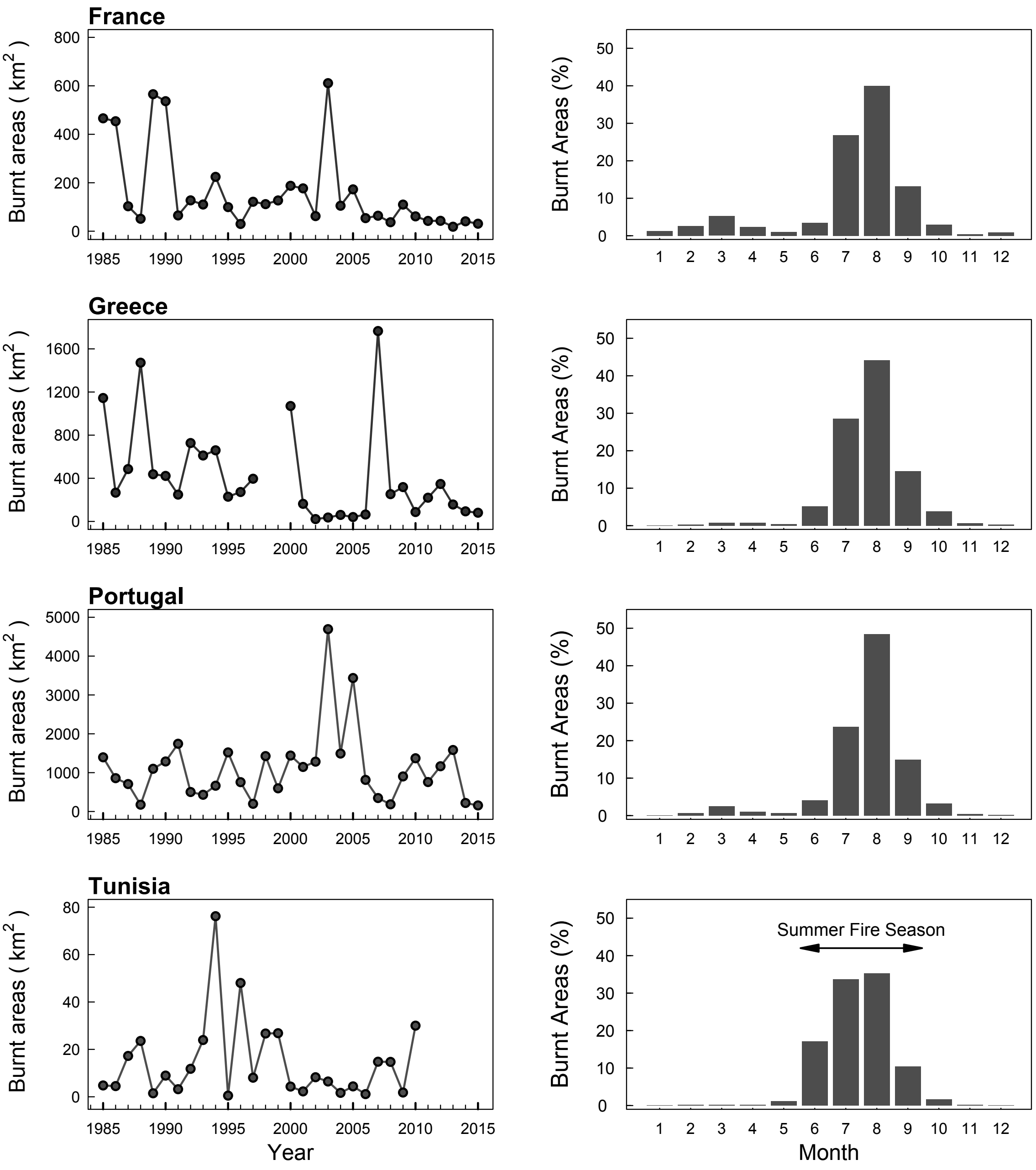


**Fig. S2** Interannual and Seasonal variability in wildfire activity for each of the four countries examined. Data is missing for years 1998 and 1999 for Greece and for the period from 2011 to 2015 for Tunisia (see details in methods). The summer wildfire season (June, July, August and September) is indicated.

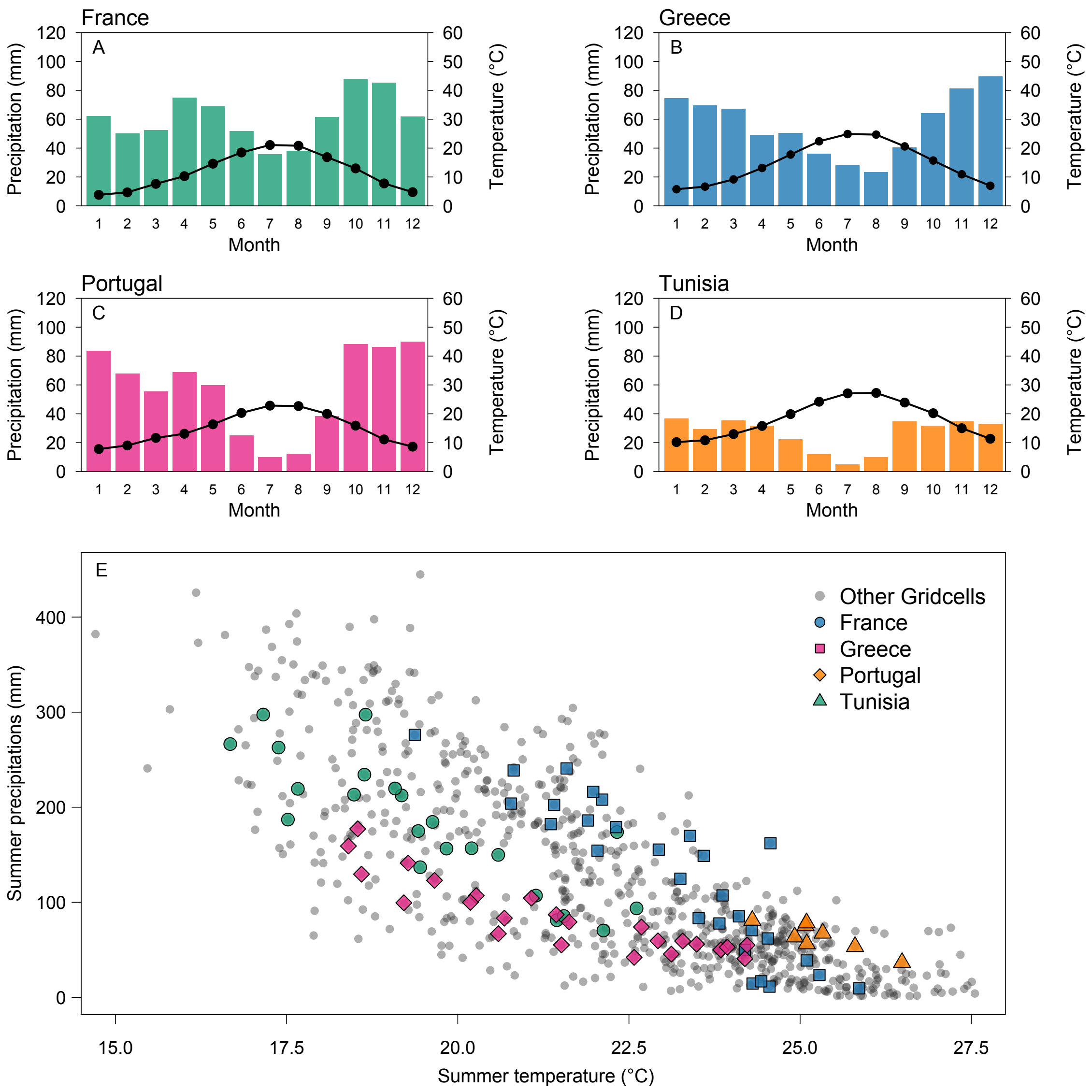


**Fig. S3** Mean climatic conditions (1985-2015) in the four countries examined (A, B, C and D) and their comparison to the rest of the Mediterranean basin (E). The four ombrothermic diagrams in A, B, C and D show the differences in summer conditions between the four countries. Subpanel E shows that the four countries examined encompasses most of the variability in mean summer conditions found in the Mediterranean Basin.

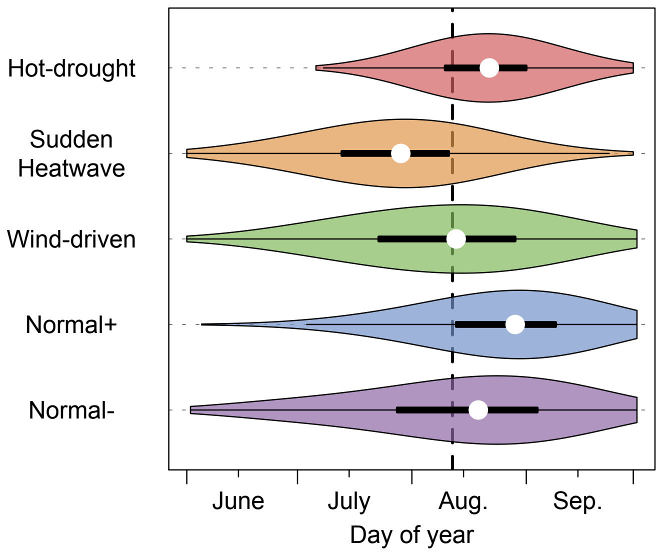


**Fig. S4** Seasonality of wildfires (>30 ha) according to each fire weather type (FWT). Violin plots show a kernel density estimation of the distribution of the day of year. White dots indicate the median of the parameter set, black boxes indicate the interquartile range between the 25th and 75th percentile, the thin black lines indicate the upper and lower adjacent values. Dashed black vertical lines denote the median for all fires. Medians for all FWTs are statistically different from each other ($p<0.001$, Pairwise Wilcoxon tests with Bonferroni correction).

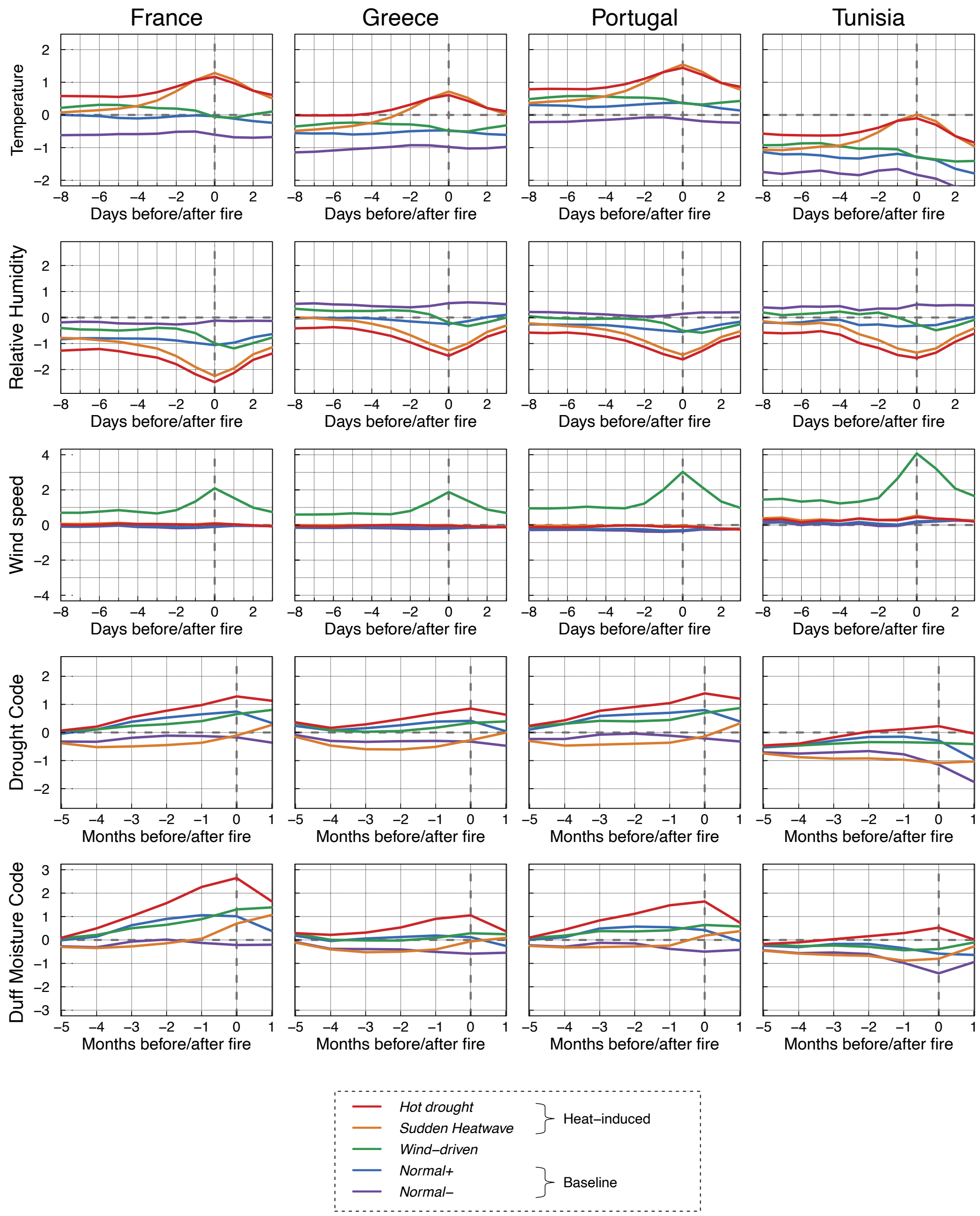


**Fig. S5** Composites of standardized anomalies associated with the different fire weather types (FWTs) relative to reference climates for each of the four countries examined. Lead lag Composites of standardized anomalies at daily (for temperature, mean relative humidity and mean wind speed) and monthly (drought code and duff moisture code) time scale are indicated. Dashed grey vertical lines denote the day (or month) of fire. The reference climates were determined by randomly selecting 10,000 non-fire gridcell*day voxels with the same location in space and time (in the annual cycle) as the fires.

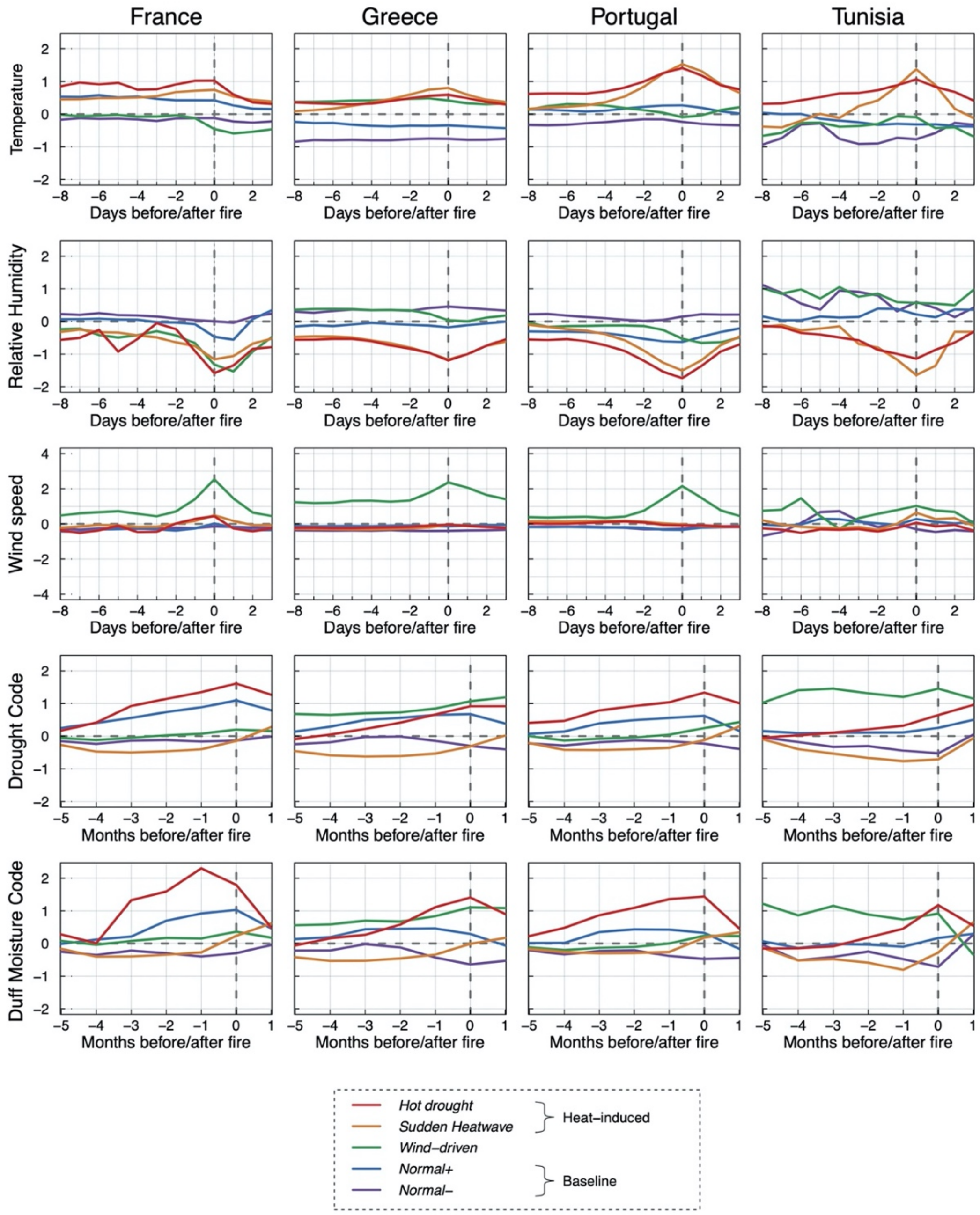


**Fig. S6** Same as Fig. S5 but composites were determined for each country only with the wildfires that occurred within that country (not the entire wildfire dataset).

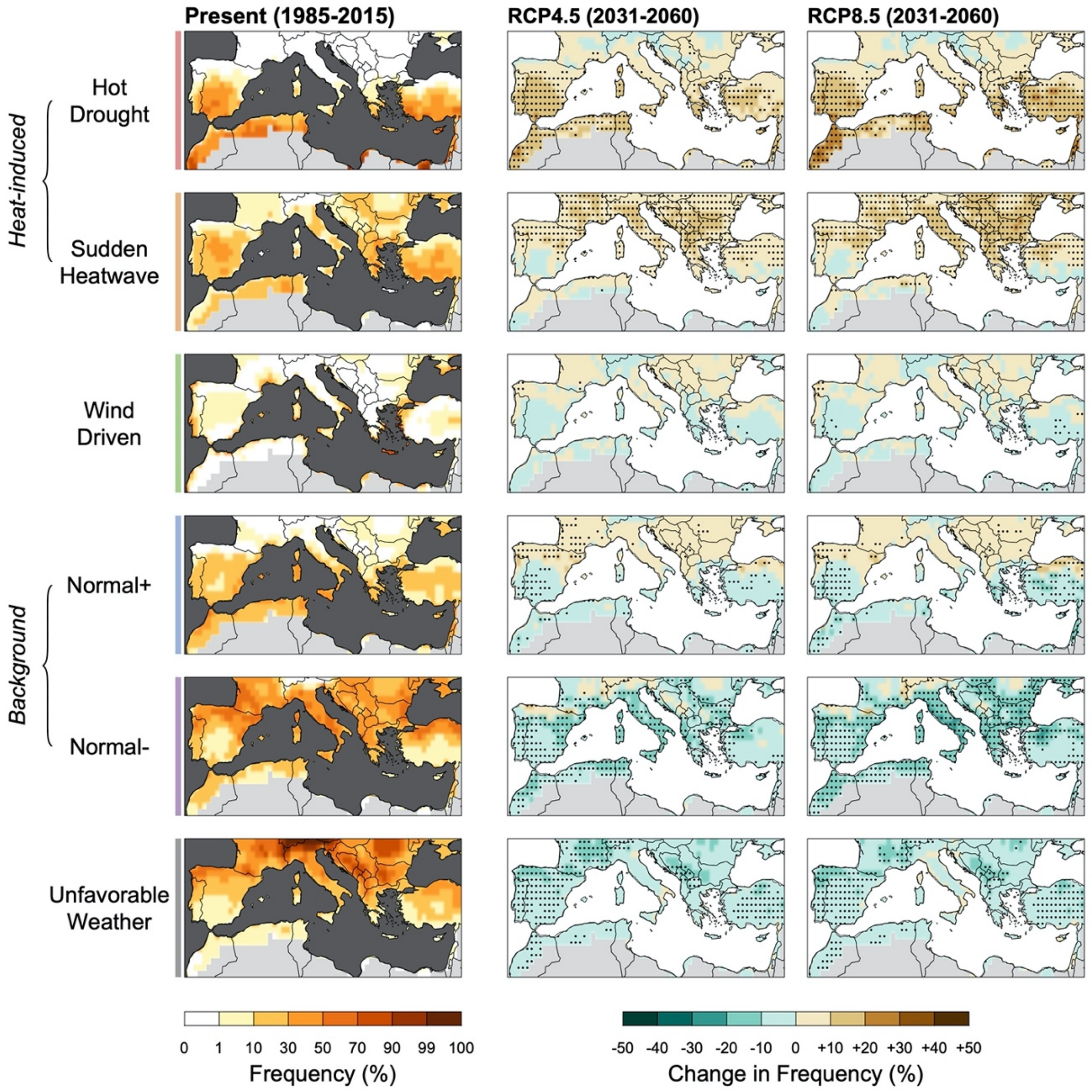


**Fig. S7** As in Fig. 3 but for the period from 2031 to 2060

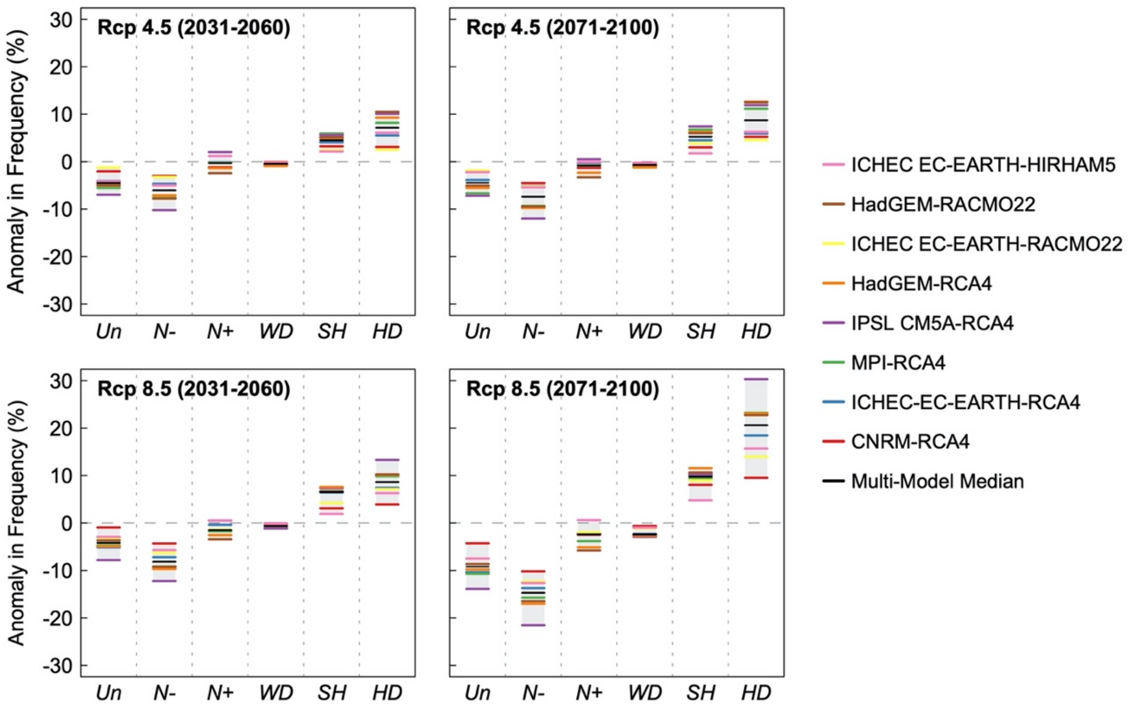


**Fig. S8** Changes in the summer frequency of fire weather types (FWTs) between the future (2031-2060; 2071-2100) and present (1985-2015) periods for two scenarios (RCP4.5 and RCP8.5) averaged over the Mediterranean Basin. The 8 GCM-RCM examined (colored dashes) and the multi-model median (black dashes) are indicated. Dashed grey horizontal lines denote zero change in frequency. *Unfavourable* (Un) *Normal-* (N-), *Normal+* (N+), *Wind-Driven* (WD), *Sudden-Heatwave* (SH) and *Hot-drought* (HD).

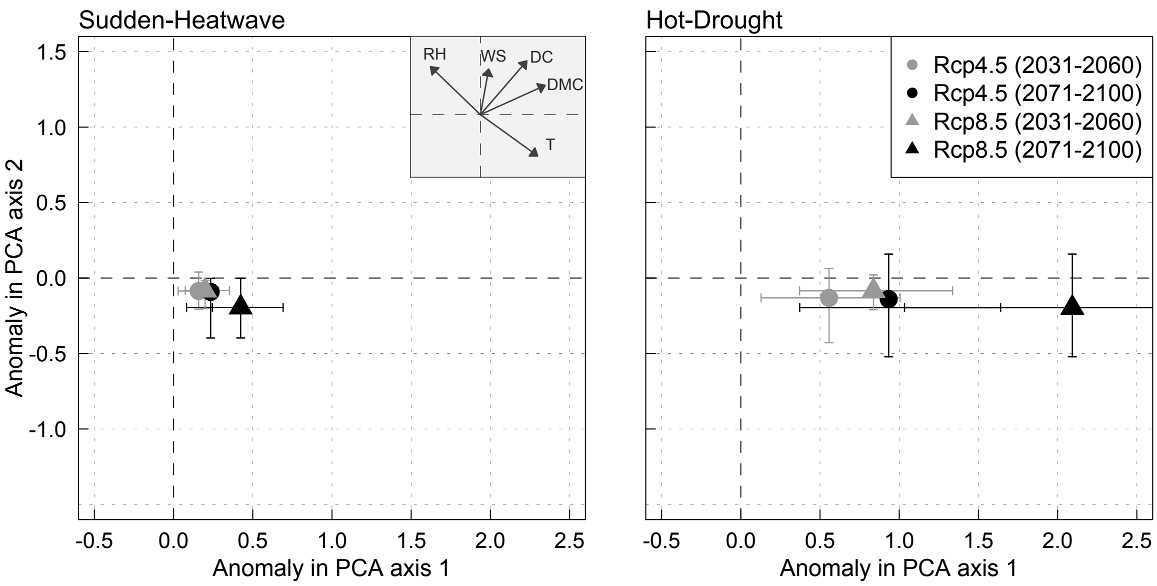


**Fig. S9** Multi model median change in the intensity of *Sudden Heatwave* and *Hot-drought* FWTs over the Mediterranean Basin shown in terms of the first two orthogonal combinations of wildfire predictors determined by principal component analysis (PCA) of five daily climate variables driving wildfires. FWT intensity is determined as the centroid of the furthest 10% gridcells*days from the FWT centroid for the entire Mediterranean Basin. Changes are expressed at end of the twenty-first century (2071–2100) relative to the present period (1985-2015) for two scenarios (RCP4.5 and RCP 8.5). Dashed black horizontal lines denote zero change in frequency. Minimum and maximum values for each of the 8 RCM models are indicated by horizontal and vertical arrows. The subpanel on the right panels indicates the loadings of climate variables in the PCA: relative humidity (RH), wind speed (WS), drought code (DC), duff moisture code (DMC) and temperature (T).

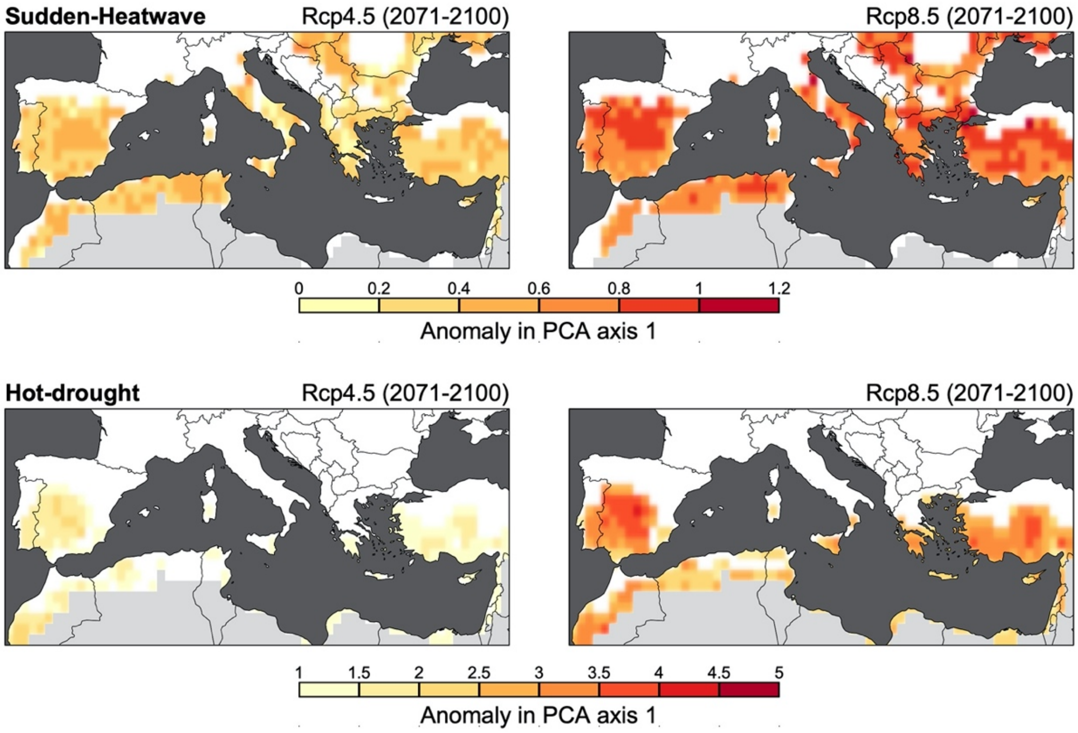


**Fig. S10** Multi model median change in the intensity of *Sudden Heatwave* and *Hot-drought* FWTs shown in terms of the first combination of wildfire predictors determined by principal component analysis (PCA) of five daily climate variables driving wildfires. FWT intensity is determined as the centroid of the furthest 10% gridcells*days from the FWT centroid for the entire Mediterranean Basin. Changes are expressed at end of the twenty-first century (2071–2099) relative to the present period (1985-2015) for two forcing scenarios (RCP4.5 and RCP8.5).

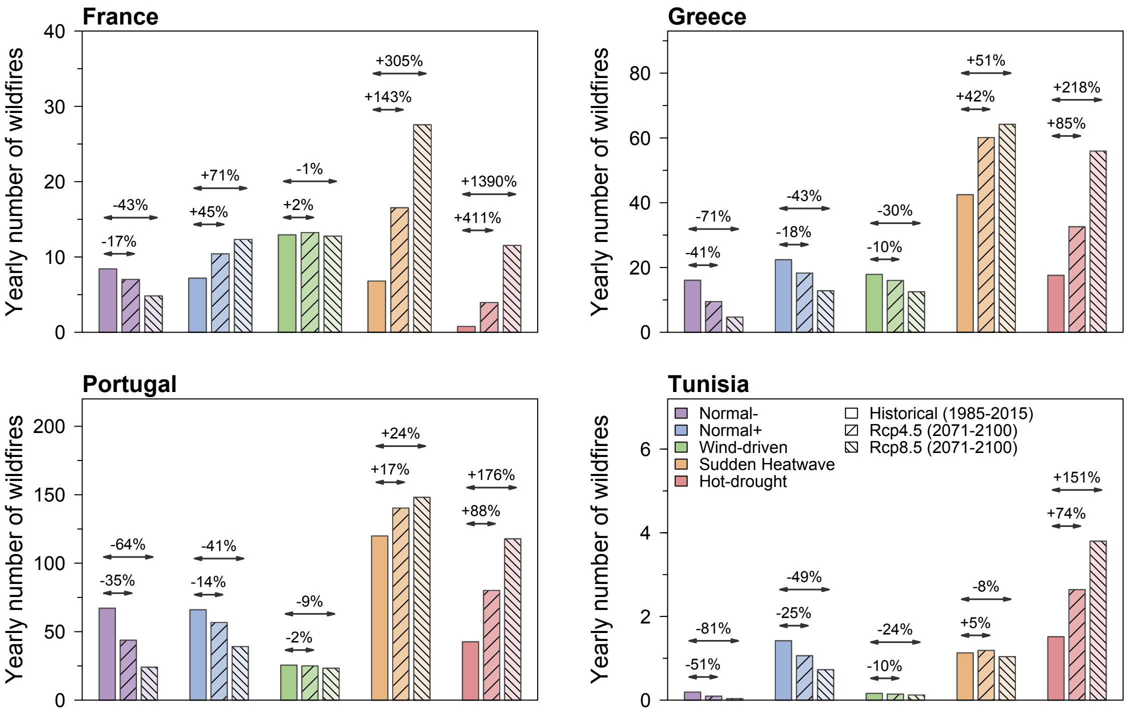


**Fig. S11** Observed and projected multi-model median of the yearly number of wildfires (>30 ha) for each fire weather type (FWT) in each of the four countries. Present (1985-2015) and future (2071-2100) periods for two scenarios (RCP4.5 and RCP8.5) are indicated.

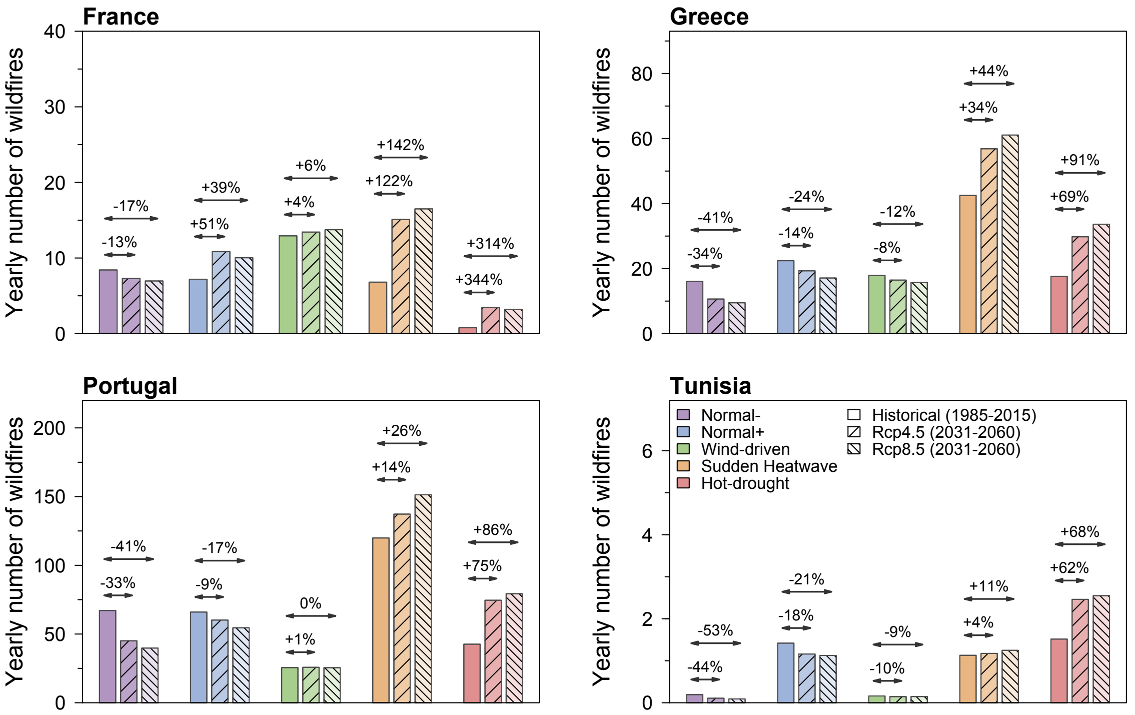


**Fig. S12** as Fig. S11 but for the period 2031-2060.

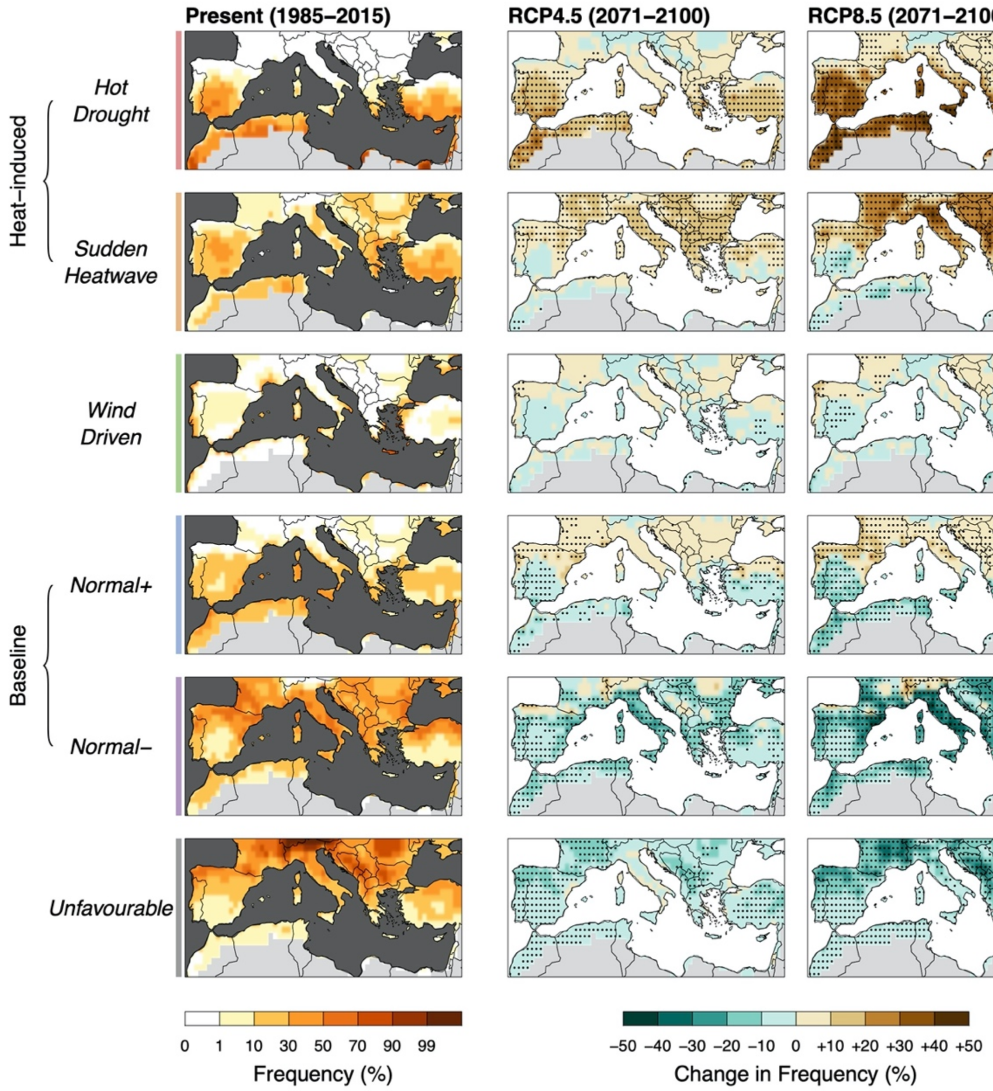


**Fig. S13** as Fig. 3 but using multivariate bias correction procedures

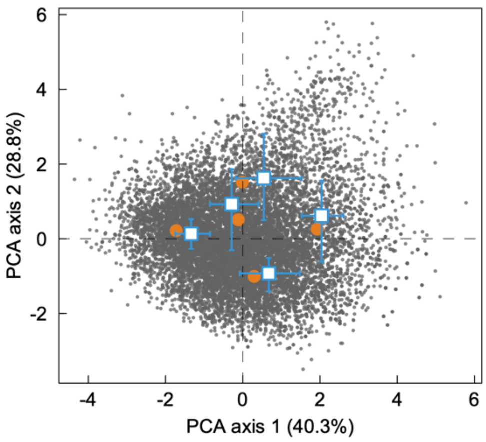


**Fig. S14** Stability of the clusters against variations in the proportion of wildfires from each country shown in terms of the first two orthogonal combinations of wildfire predictors determined by principal component analysis (PCA). Blue squares indicate the medians of centroids resulting from the k-means analysis performed on 1,000 bootstrap-resampled wildfire datasets, in which the number of wildfires from each country was proportional to the corresponding number of fire gridcells. Error bars show the 5 and 95% confidence intervals. Daily values of the climate variables associated with each wildfire (grey dots) and centroids (orange dots) resulting from the original wildfire dataset (as in Fig. 1a.) are indicated for reference.
